# The Effects Of Heavy Metals On The Morphological And Biochemical Composition Of The Blood Of Young Cattle Taking Into Account Of Breed Features

**DOI:** 10.1101/2020.08.02.232694

**Authors:** Rustem B. Temiraev, Yusup A. Yuldashbaev, Susanna K. Cherchesova, Sofia F. Lamarton, Valentina S. Gappoeva, Valery R. Kairov, Alja N. Doeva, Arthur L. Kalabekov, Inna I. Kornoukhova

## Abstract

Heavy metals can selectively accumulate in certain organs and remain in them for a long time. As a result, the accumulation of metal in one or another organ may cause strongest intoxication of the animal. The purpose of the research is to study the morphological and biochemical composition of the blood of young cattle of different breeds of the dairy and dairy-and-beef direction of productivity fed in the technogenic zone in the RNO-Alania diets with a high content of heavy metals. According to the principle of analogues, taking into account the breed, origin, age, sex and body weight, 4 groups of 10 animals each of different breed were formed: Group I - black and white, Group II - Danish Red, Group III - Swiss, and Group IV - Simmental. The experimental material was statistically processed using the Microsoft Excel software for mathematical analysis. With a tendency of unreliable differences in morphological indices of blood, in diminishing numbers of erythrocytes and hemoglobin, the test breeds should be arranged in the following order: Simmental → Swiss → Red Danish → Black and White. At the same time, as compared to the animals of the black-and-white breed, the young cattle of the Simmental breed showed the best level of intermediate metabolism, which resulted in an increase in total protein and sugar in the blood, which indicates an improvement in their protein and energy metabolism; an increase in the content of albumin in the liquid internal medium and γ-globulins;an increase inconcentration oftotal lipids in serum, respectively, while reducing the level of cholesterol, which indicates an improvement fat metabolism in animals; a significant (P <0.05) decrease in blood saturation with cadmium, lead and zinc, respectively, by 26.8, 48.8 and 30.6%. The highest content of these elements was noted in the wool, then in decreasing order: in bone tissue → liver → muscle tissue → lungs → kidneys. Moreover, between the level of heavy metals in muscles, liver and lungs, on the one hand, and in the wool, bone tissue and kidneys, on the other hand, there was an inversely proportional relationship.

## Original Articles

Articles that present a contribution which is entirely new to knowledge and allow other researchers, based on the written text, to judge the conclusions, check the accuracy of the analyzes and deductions of the author and repeat the investigation if they so wish.

## Relevance of the topic

The most important task of increasing the productivity of young fattening cattle is the development of flexible technologies for the production of beef, aimed at the maximum and rational use of their own feed, natural pastures with minimal consumption of concentrated feed. A promising direction of stabilization and development of beef cattle breeding in Russia should be considered the intensification of production based on modern scientific achievements, new technologies that ensure high productionperformance of animals and labor productivity, environmental friendliness and competitiveness [1, 2, 3].

Heavy metals have a high capacity for diverse chemical and biochemical reactions. Many of them have variable valence and are involved in redox processes. Heavy metals and their compounds, like other chemical compounds, are able to move and redistribute in living environments, i.e. migrate. As a result of interconversions between the ingested metals or their compounds and the chemicals of various tissues and organs, new metal compounds canform, which have different properties and behave differently in the body. These toxicants can selectively accumulate in certain organs and remain in them for a long time. As a result, the accumulation of metal in one or another organ may be a manifestation of the strongest intoxication of the animal’s body [4, 5, 6, 7].

The Republic of North Ossetia – Alania (RNO-Alania) is one of the most heavily polluted areas in Russia due to the high concentration of industrial enterprises in the city of Vladikavkaz. The main environmental polluters are enterprises of non-ferrous metallurgy, JSC Magnit, JSC Electrozinc, JSC Pobedit and others. Among heavy metals, mercury, lead, zinc and cadmium lead a number of pollutants due to their high rates of technogenic accumulation in the environment. The maximum number of the first three elements was found in the city of Vladikavkaz and in a radius of up to 15 km, which also includes the territory of the Prigorodny district of North Ossetia – Alania [8, 9, 10].

The range of environmental effects at the molecular, tissue, and cellular levels largely depends on the concentration and duration of exposure of the elements, its combination with other factors, the preceding state of health of the animal and its immunological reactivity. Of great importance is the genetically determined sensitivity to the influence of certain heavy metals [11, 12, 13].

On this basis, in the conditions of the technogenic zone of North Ossetia - Alania, the problem of studying the influence of heavy metals on the state of intermediate metabolism of fattened young cattle of the dairy and dairy-and-beef direction of productivity, taking into account pedigree characteristics, is highly relevant.

**The purpose of the research** is to study the morphological and biochemical composition of the blood of young cattle of different breeds of the dairy and dairy-and-beef productivity fed diets with a high content of heavy metals in the technogenic zone in the RNO – Alania.

## Material and research methods

To achieve this goal, in the conditions of the farm of “Meat Products” RNO – Alania, research and commercial experiment was conducted. The objects of the research were young fattening bulls of black-and-white, red Danish, Swiss and Simmental breeds, that is, young dairy and dairy-and-beef breeds bred in our region.

According to the research scheme (Table 1), 40 bulls at the age of 6 months were selected, of which 4 groups of 10 animals each of different breed were formed according to the principle of analogues, taking into account the breed, origin, age, sex and body weight: I group - black and white, II group - red Danish, III group - Swiss and IV - Simmental.

**Table 1.**
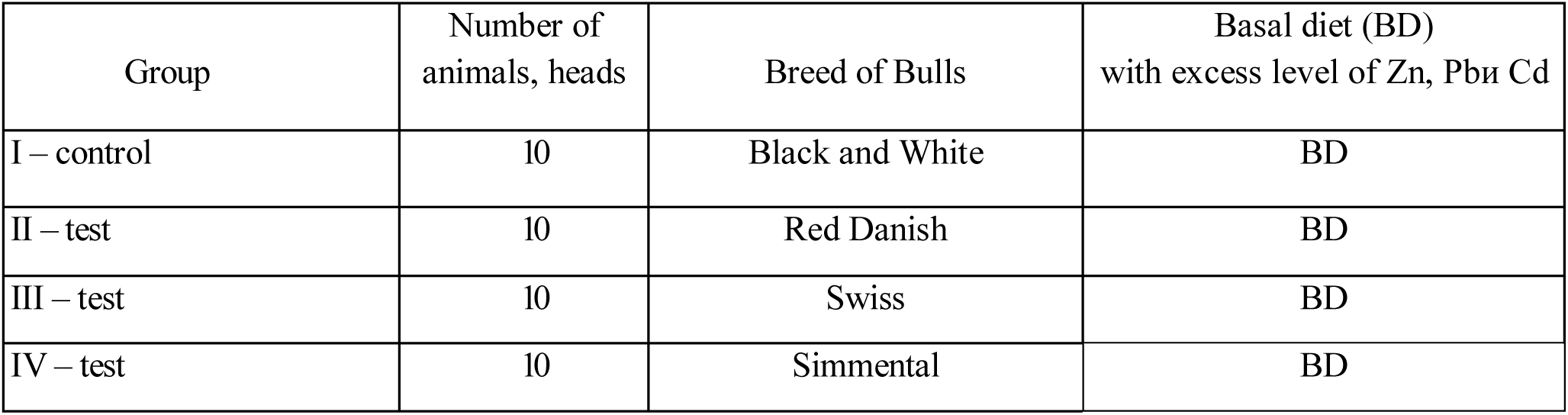
Scheme of the research and commercial experiment.

The conditions for feeding and keeping the experimental animals were similar. The animals of the compared groups were fed with rations balanced in accordance with detailed norms of the RAAS [14].

In the course of the research, average samples of the entire range of feed used in feeding experimental animals were taken monthly, which were subsequently subjected to full chemical analysis according to generally accepted methods [15]. The content of heavy metals (Cd, Pb, Zn) in samples of feed and blood of experimental animals was determined by the atomic adsorption method on an AAZ-115-M1 spectrophotometer.

Heavy metals, as bivalent elements, can inhibit the microflora of ruminant forestomach and inhibit the activity of enzymes involved in the hydrolysis of organic polymers of feed. With this in mind, the rations of animals of all groups included the enzyme preparation Celloviridin G20x at a dose of 0.01% of the dry matter norm.

The conditions of keeping bulls of the compared breeds were the same. The young fattening cattle werekept in stables.

To study the morphological and biochemical parameters in experimental animals (3 heads from a group) at the age of 6 and 18 months, blood was taken from the jugular vein in the morning before feeding, which was stabilized with heparin. In the blood of experimental animals according to generally accepted methods [16], indicators of the morphological and biochemical composition of the blood were studied.

The experimental material was statistically processed using the Microsoft Excel software for mathematical analysis.

## Research results and discussion

The territory of the Prigorodny District, where the RNO-Alania Meat Products Farm is located, is an area with a high content of heavy metals in the soil and forage crops, which is caused by emissions from large non-ferrous metallurgy enterprises located in the city of Vladikavkaz.

The composition and nutritional value of the rations of experimental animals varied depending on the age period: 6–9 months, 9–12 months, 12–15 months, and 15–18 months.

In the course of a research and production experiment, we studied the content of heavy metals in daily rations of young animals of compared breeds, depending on the age period. In the diets of the experimental bulls of the compared groups there was an excess of the zinc rate, respectively: at the age of 6– 9 months - 2.14 times; 9-12 months - 2.48; at the age of 12-15 months - 1.90 and at the age of 15-18 months - 1.90 times. In these age periods in the rations of experimental bulls, the amount of lead was 124.26; 192.46; 216.96 and 264.00 mg and cadmium - 10.27; 13.27; 14.91 and 18.18 mg, respectively.

Despite the constant flow into the blood of ruminants and the release of various nutrients from it, the composition of blood components is highly stable due to the regulation of metabolic processes of the nervous and humoral systems. But all the same, some blood parameters, depending on the conditions of feeding may vary in one direction or another.

Based on the foregoing, we studied the dynamics of the morphological indices of the blood of animals of the compared breeds with an excessive supply of heavy metals with feed (Table 2).

**Table 2.**
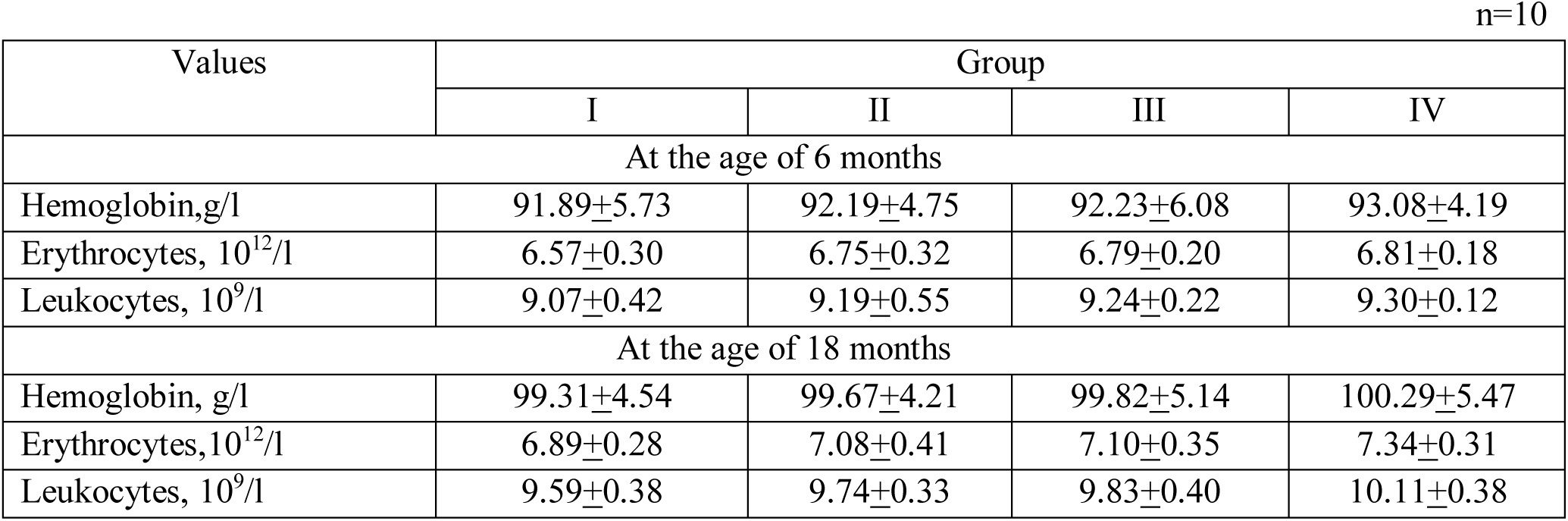
Dynamics of morphological values inblood of experimental animals.

The morphological parameters of blood in animals of the compared groups with age and under the influence of breed characteristics changed in accordance with biological laws. Moreover, the increase in the intensity of growth under the influence of age features was accompanied by a tendency to increase the absolute data characterizing the number of erythrocytes, hemoglobin and leukocytes in experimental animals.

Of the morphological parameters of blood, erythrocytes and hemoglobin were the most susceptible to changes under the action of the alimentary factor. It has been established that the amount of hemoglobin and erythrocytes that perform the most important functions of supplying organs and tissues with oxygen, adsorption and delivery of free amino acids contained in the blood, had a direct biological connection with the growth rate of animals of comparable breeds, which is associated with the metabolic rate in the body. Therefore, the blood of the bulls of the Simmental breed had the highest content of erythrocytes and hemoglobin, which surpassed the control by these parameters by 0.45 × 10^12^/l (P>0.05) and - by 0.98 g/l (P>0.05) respectively, and according to their decreasing quantity, the test breeds should be arranged in the following order: Swiss → Red Danish → Black-and-White.

Leukocytes in the body of animals perform protective functions. In the course of the experiment, it was found that there were practically no differences in the number of leukocytes in the blood of the compared groups of bulls.

Serum proteins perform a protective function, transport fats, vitamins and metabolic products. Therefore, to study the effect of breed characteristics on protein metabolism in the body of bulls at the age of 18 months, the level of total protein and its fractions in the blood serum was determined (Table 3).

**Table 3.**
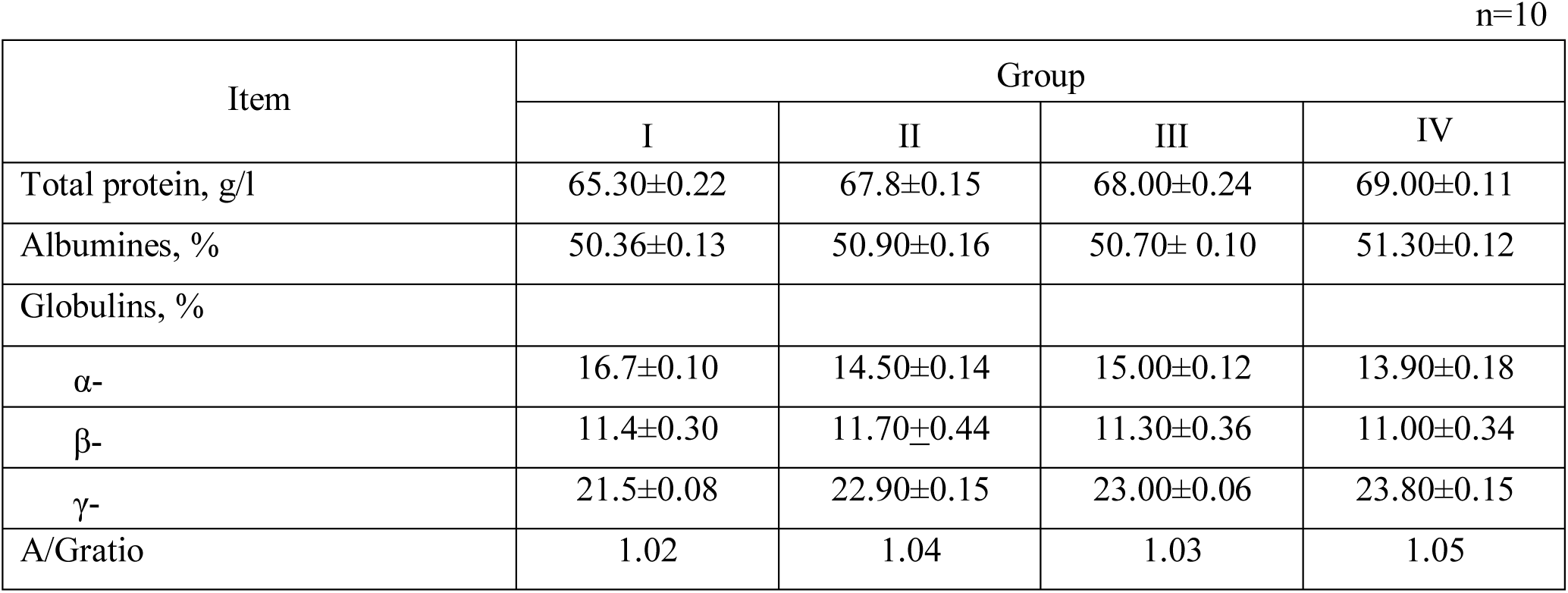
The content of protein and protein fractions in the blood serum of bulls.

It was established that the concentration of total protein in the blood serum had a directly proportional relationship with the level of nitrogen utilization of feed and the growth energy of the experimental bulls. Therefore, the lowest level of serum proteins was in the blood of animals of group I - 65.3 g/l, and the highest - in the blood of young animals in group IV - 69.0 g/l, which is 3.7 g/l (P <0.05) more than in the control. This indicates a better protein metabolism in Simmental animals compared to the young stock of other breeds.

One of the main indicators of metabolic tension in young ruminants is the total protein content in blood serum. The study of the concentration of protein fractions in the blood, as well as the A/G protein ratio, was important for determining the effect of breed characteristics on the physiological state and resistance of the organism.

In the course of the experiment, animals of group IV surpassed the peers of group I in the albumin content by 0.94% (P <0.05). With an excessive content of heavy metals in animal feed in specialized meat breeds, the protective functions of the body were higher. Thus, in relation to the peers of the first experimental group in animals of the fourth experimental group, there was an increase in the number of γ- globulins in blood serum by 2.3% (P <0.05).

Due to the increase in the blood serum of animals of the Simmental breed, the number of albumins and, especially, γ-globulins, the A / G ratio was higher compared with the peers of group I by 0.03 units.

Alpha- and beta-globulins, along with albumins, play a large role in the transport of nutrients and have enzymatic activity. The content of these globulin fractions in the blood serum of the bulls of the compared breeds was within the physiological norm and there were no significant differences between the animals of the compared groups.

The most important qualitative characteristic of the influence of heavy metals on the intermediate metabolism is the biochemical composition of the blood of experimental animals (Table 4).

**Table 4.** Dynamics of biochemical values of blood serum of experimental bulls.

| Values | Group |  |  |  |
| --- | --- | --- | --- | --- |
|  | I | II | III | IV |

| At the age of 6 months |  |  |  |  |
| --- | --- | --- | --- | --- |
| Sugar, mmol/l | 2.41±0.02 | 2.40±0.03 | 2.40±0.04 | 2.41±0.03 |
| Total lipids, mmol/l | 1.80±0.02 | 2.20±0.04 | 2.22±0.03 | 2.28±0.01 |
| Calcium, mmol/l | 2.48±0.12 | 2.49±0.18 | 2.49±0.13 | 2.48±0.15 |
| Phosphorus, mmol/l | 1.54±0.12 | 1.57±0.10 | 1.56±0.10 | 1.57±0.12 |
| At the age of 18 months |  |  |  |  |
| Sugar, mmol/l | 3.61±0.02 | 3.68±0.04 | 3.83±0.03 | 3.91±0.05 |
| Total lipids, mmol/l | 4.49±0.11 | 4.55±0.08 | 4.63±0.05 | 4.72±0.09 |
| Calcium, mmol/l | 2.84±0.18 | 3.11±0.32 | 3.29 ±0.22 | 3.30±0.11 |
| Phosphorus, mmol/l | 1.88±0.09 | 2.06±0.03 | 2.17± 0.06 | 2.25±0.08 |

At the age of 18 months there was an increase in the concentration of sugar in the blood of bulls of experimental groups. At the same time, the highest level of blood sugar was found in the young animals of the IV experimental group - 3.91 mmol/l, which is 0.30 mmol/l (P <0.05) more than in the control.

The content of total lipids in the bulls of the compared groups at the age of 6 months in the blood serum is markedly lower than at 18 months of age, which is due to puberty. By concentration of total lipids in blood serum, bulls of the IV and III groups significantly (P <0.05) surpassed theanimals of group II, respectively, by 0.17 mmol/l and by 0.08 mmol/l, experimental animals of group I, respectively - by 0.23 and 0.14 mmol/l. This indicates the best lipid metabolism in Simmental animals.

It is known that the ratio of calcium and phosphorus in the blood of animals depends not only on their quantity, but also on the ratio of elements in the fed diet.

Of the mineral elements, calcium and phosphorus are found in the body of animals in the largest amounts. The content of total calcium and inorganic phosphorus corresponded to the physiological norm. Moreover, the content of these macroelements in serum was, respectively, 0.46 (P <0.05) and 0.37 mmol/l (P <0.05) higher in the animals of group IV in comparison with the bulls of group I.

When assessing the metabolism under conditions of excessive content of heavy metals in feed, it was important to assess the effect of the breed characteristics of fattened young cattle on the ecological characteristics of their internal liquid medium. Therefore, in the course of the research we also studied the content of zinc, lead and cadmium ions in the blood serum of fattened young cattle of the compared breeds (Table 5).

**Table 5.**
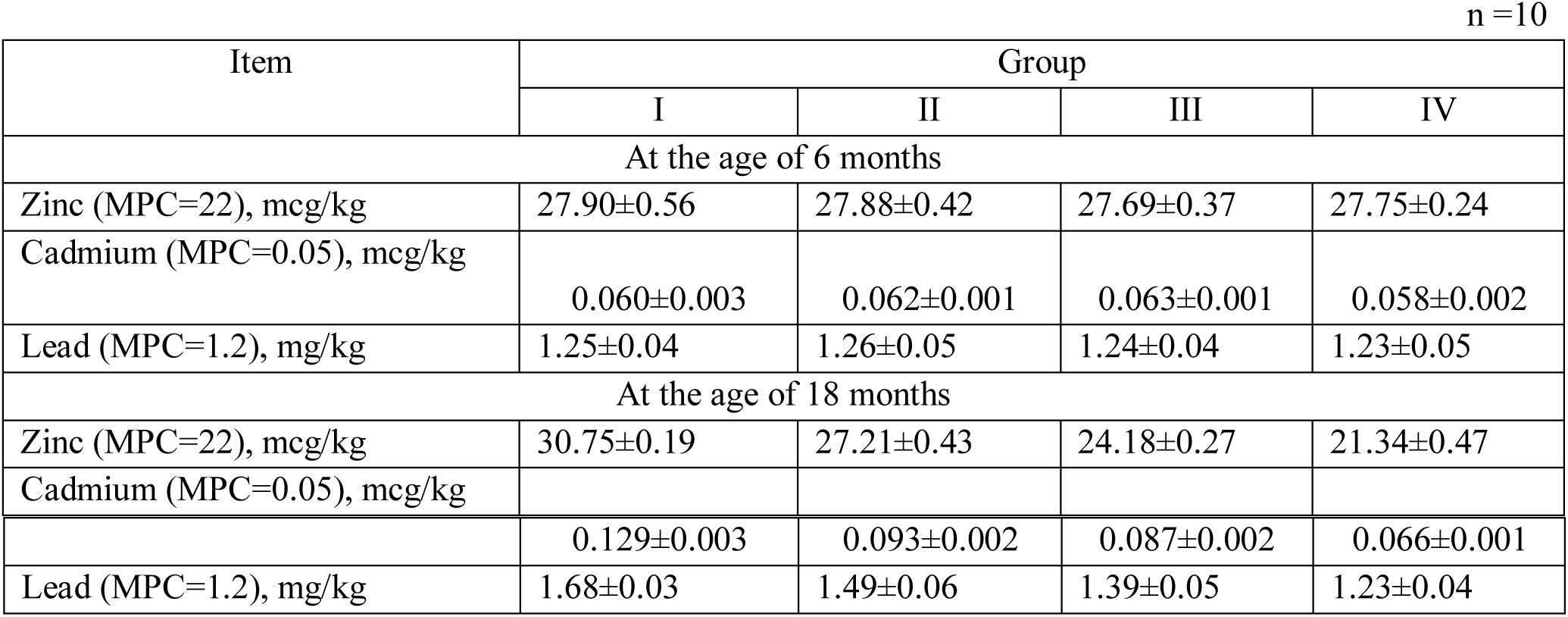
The content of heavy metals in the blood of experimental bulls.

| Item | Group |  |  |  |
| --- | --- | --- | --- | --- |
|  | I | II | III | IV |
| At the age of 6 months |  |  |  |  |
| Zinc (MPC=22), mcg/kg | 27.90±0.56 | 27.88±0.42 | 27.69±0.37 | 27.75±0.24 |
| Cadmium (MPC=0.05), mcg/kg | 0.060±0.003 | 0.062±0.001 | 0.063±0.001 | 0.058±0.002 |
| Lead (MPC=1.2), mg/kg | 1.25±0.04 | 1.26±0.05 | 1.24±0.04 | 1.23±0.05 |
| At the age of 18 months |  |  |  |  |
| Zinc (MPC=22), mcg/kg | 30.75±0.19 | 27.21±0.43 | 24.18±0.27 | 21.34±0.47 |
| Cadmium (MPC=0.05), mcg/kg |  |  |  |  |
| n =10 |  |  |  |  |
|  | 0.129±0.003 | 0.093±0.002 | 0.087±0.002 | 0.066±0.001 |
| Lead (MPC=1.2), mg/kg | 1.68±0.03 | 1.49±0.06 | 1.39±0.05 | 1.23±0.04 |

Heavy metals are lipophilic, so their serum level was directly proportional to the concentration of total lipids. The content of heavy metals in the blood of animals of all tested breeds was higher than the maximum permissible level (MPC). However, at the age of 6 months, their serum concentration was approximately uniform.

The highest concentration of cadmium is 0.129 mg/kg, lead - 1.68 mg/kg and zinc - 30.75 mg/kg was observed at 18 months of age in the blood serum of young animals of I group (black-and-white breed), and the lowest – in animals of group IV (Simmental breed, reliably (P <0.05) inferior to them in blood saturation with these toxicants, respectively, by 26.8, 48.8 and 30.6%. Moreover, it should be noted that although the concentration of zinc, lead and cadmium in the serum of young Simmental breed exceeded the MPC, but their level was close to the lowestpoint of the maximum permissible level.

With this in mind, we studied the distribution of these toxicants in the organs and tissues of bulls of various breeds fed rations enriched with Cellobacterin G20x (Table 6).

**Table 6.** The content of heavy metals in organs and tissues of young bulls.

| Organs and tissues | Group |  |  |  |
| --- | --- | --- | --- | --- |
|  | I | II | III | IV |
| Zinc content, mg/kg |  |  |  |  |
| Muscular | 74.13±0.15 | 64.56±0.18 | 56.31±0.22 | 39.07±0.11 |
| Bony | 119.8±0.29 | 131.8±0.31 | 182.5±0.27 | 221.8±0.39 |
| Liver | 91.8±0.16 | 88.8±0.17 | 69.5±0.21 | 46.8±0.18 |
| Kidney | 41.14±0.17 | 48.55±0.12 | 68.78±0.11 | 78.11±0.15 |
| Lungs | 71.44±0.21 | 67.51±0.14 | 52.35±0.20 | 41.07±0.16 |
| Wool | 138.9±0.24 | 151.8±0.31 | 262.5±0.37 | 279.8±0.41 |
| Lead content, mg/kg |  |  |  |  |
| Muscular | 0.62±0.002 | 0.55±0.001 | 0.49±0.002 | 0.37±0.003 |
| Bony | 1.98±0.004 | 2.19±0.003 | 2.61±0.002 | 3.42±0.005 |
| Liver | 0.98±0.003 | 0.93±0.0030 | 0.63±0.003 | 0.53±0.003 |
| Kidney | 0.26±0.003 | 0.29±0.002 | 0.38±0.003 | 0.48±0.004 |
| Lungs | 0.60±0.001 | 0.56±0.001 | 0.32±0.002 | 0.29±0.002 |
| Wool | 2.18±0.003 | 2.30±0.0023 | 3.56±0.004 | 3.68±0.004 |
| Cadmium content, mg/kg |  |  |  |  |
| Muscular | 0.071±0.001 | 0.065±0.001 | 0.059±0.001 | 0.044±0.001 |
| Bony | 0.289±0.002 | 0.314±0.002 | 0.378±0.001 | 0.443±0.002 |
| Liver | 0.097±0.001 | 0.089±0.001 | 0.062±0.001 | 0.055±0.001 |
| Kidney | 0.027±0.003 | 0.032±0.003 | 0.039±0.003 | 0.047±0.003 |
| Lungs | 0.061±0.001 | 0.057±0.001 | 0.036±0.002 | 0.030±0.002 |
| Wool | 0.297±0.002 | 0.318±0.002 | 0.408±0.004 | 0.459±0.003 |

Based on the foregoing, the animals of the compared breeds, taking into account the accumulation of heavy metals in the listed organs and tissues, can be arranged in a decreasing order in the following order: Simmental → Swiss → Red Danish → Black-and-White breeds. The highest content of heavy metals was in the muscles, liver and lungs of the young animals of group I (black-and-white breed);it was significantly (P <0.05) ahead the animals of group IV (Simmental breed) by the concentration of zinc in them 1.90, 1, 96 and 1.74 times, lead - by 1.68, 1.85 and 2.07 times and cadmium - by 1.61, 1.76 and 2.03 times respectively. Moreover, in young Simmental breed the concentration of these elements in no case exceeded the MAC.

There was an inversely proportional relationship between the level of heavy metals in muscles, liver, and lungs, on the one hand, and in wool, bone tissue, and kidneys, on the other hand. Therefore, the highest content of these toxicants in wool, bone tissue and kidney showed young bulls of group IV, significantly (P <0.05) being ahead of animals of group I by zinc concentration – 2.01, 1.85 and 1.90 times, lead - 1.69, 1.73 and 1.85 times and cadmium - 1.54, 1.53 and 1.74 times respectively. In our opinion, the reasons for this are, on the one hand, the fact that bone tissue and hair cover are the main depot of toxicants, on the other hand, the fact that, through the kidneys, more zinc, lead and cadmium were removed with the urine in animals of the Simmental breed than in other breeds of their peers. The last one point is consistent with the data on balance of heavy metals in the test animals during the physiological experiment.

## Conclusion

1. With a tendency of unreliable differences in morphological blood values, according to diminishing numbers of erythrocytes and hemoglobin, the tested breeds should be arranged in the following order: Simmental → Swiss → Red Danish → Black-and-White. At the same time, in relation to animals of the black-and-white breed, young animals of the Simmental breed were characterizedby better level of intermediate metabolism, which expressed itself:
  - in a reliable (P <0.05) increase in blood of total protein by 3.7 g/l and sugar - by 0.30 mmol/l, which indicates an improvement in their protein and energy metabolism;
  - in a significant (P <0.05) increase in the content of albumin in the liquid internal medium by 0.94% and γ-globulins - by 2.3%;
  - in a significant (P <0.05) increase of concentration of total lipids in the blood serum, respectively, by 0.23 mmol/l while reductionof the cholesterol level, which indicates an improvement in their fat metabolism;
  - in a reliable (P <0.05) increase in blood of the concentration of calcium and phosphorus by 0.46 and 0.37 mmol/l 44.2%;
  - in a reliable (P <0.05) decrease in blood saturation with cadmium, lead and zinc, respectively, by 26.8, 48.8 and 30.6%.
2. The concentration of zinc, lead and cadmium in the organs and tissues of young buuls had significant fluctuations. The highest content of these elements was foun in wool, then in decreasing order: in bone tissue → liver → muscle tissue → lungs → kidneys. Moreover, between the level of heavy metals in muscles, liver and lungs, on the one hand, and in wool, bone tissue and kidneys, on the other hand, there was an inversely proportional relationship. At the same time, it has been found that with respect to the control:
  - the animals of the Simmental breed had significantly (P <0.05) less zinc concentration in their muscles, liver and lungs 1.90, 1.96 and 1.74 times, lead - 1.68, 1.85 and 2.07 times and cadmium - 1.61, 1.76 and 2.03 times respectively. Moreover, in young Simmental breed the concentration of these elements in no case exceeded the MAC.
  - young bulls of group IV differed significantly (P <0.05) with a higherzinc level in wool, bone and kidney 2.01, 1.85 and 1.90 times, lead - 1.69, 1.73 and 1, 85 times and cadmium - 1.54, 1.53 and 1.74 times respectively.

